# Comparative genomics of epiphytic and endophytic bacterial culture collections from *Arabidopsis thaliana* in Ōtautahi (Christchurch), Aotearoa New Zealand

**DOI:** 10.64898/2025.12.14.694188

**Authors:** Moritz Miebach, Rudolf O. Schlechter, Renji Jiang, Cassidy Weaver, Simisola O. Oso, Christian Stocks, Markosovo Marjaya, Evan Kear, Sandra Hirsch, Mila Oeltjen, Luis R. Paniagua Voirol, Matthew B. Stott, Students of the MSc course “Leaf surface microbiology 2021-2025”, Mitja N.P. Remus-Emsermann

**Affiliations:** Te Kura Pūtaiao Koiora | School of Biological Sciences, Te Whare Wānanga o Waitaha | University of Canterbury, Aotearoa New Zealand; Institute of Biology -Microbiology and Dahlem Centre of Plant Sciences, Freie Universität Berlin, Berlin, Germany

**Keywords:** Bacterial culture collection, Arabidopsis thaliana, New Zealand, full genome sequencing, Phyllosphere, Microbiota, Graduate class research

## Abstract

Bacterial culture collections are essential resources for exploring the diversity of microorganisms and their interactions with each other and their hosts. Here, we report on the sequencing of the first 129 bacterial isolates, representing 34 genera, from a culture collection of more than 600 bacterial strains originally isolated from leaves of a naturalised *Arabidopsis thaliana* population from Ōtautahi (Christchurch), Aotearoa New Zealand. Epiphytic (leaf surface), and endophytic (apoplastic) bacteria were isolated separately from the same leaves, providing complementary insights into both compartments. The recovered isolates encompass the dominant taxa typically associated with the *Arabidopsis* phyllosphere, including *Pseudomonas*, *Sphingomonas*, *Methylobacterium*, and *Flavobacterium*. Their full genome assemblies (BUSCO average completeness > 99%, checkM average completeness > 97% and average contamination < 1%) were analysed and compared to assess genomic features across epiphytic and endophytic lineages. While the epiphytic and endophytic strain collections did not show large genomic differences, certain functional categories differ, such as terpene biosynthesis and biofilm formation being enriched in epiphytic strains, while arginine biosynthesis and carbohydrate degradation were associated with endophytic strains. These data provide a genomic foundation for future experimental work on leaf-associated microbial ecology and plant–microbe interactions. To our knowledge, this is the first *Arabidopsis* leaf culture collection established from a Southern Hemisphere source.

## Introduction

Plant leaves are colonised by a wide range of bacteria, with epiphytic communities inhabiting the phyllosphere (leaf surfaces) and endophytic communities residing in the apoplast (leaf interior) (Schlechter et al. 2019). These complex communities influence the fate of their host by affecting health, disease and development (Stone et al. 2018; Zhu et al. 2022). In turn, plant microbiota are determined by both the host species and its location (Knief, Ramette, et al. 2010; Karasov et al. 2024; Redford et al. 2010). Although many studies have described plant-associated microbiota, the processes governing community assembly and host selection remain poorly understood. To unravel these mechanisms, it is essential to study microbial interactions under highly controlled experimental conditions (Vorholt et al. 2017).

Gnotobiotic systems—plants grown under axenic conditions and colonised with defined microbial communities—provide a powerful framework for such mechanistic studies (Vorholt et al. 2017). The establishment of these systems depends on two factors: first, the ability to surface-sterilise seeds without compromising germination, a method which has been well established for *Arabidopsis thaliana* (Arabidopsis from here onwards) (Miebach et al. 2020; Bodenhausen et al. 2014); and second, the availability of microbial isolates suitable for community reconstitution (Vorholt et al. 2017). Thus far, there have been few published efforts to produce collections of microbial isolates from *Arabidopsis* roots and leaves suitable for these systems.

Comprehensive bacterial culture collections are often thematic “metacollections” that originate from many different sample types such as the New Zealand-based International Collection of Microorganisms from Plants (ICMP), the Belgian BCCM/LMG Bacteria Collection, the German Collection of Microorganisms and Cell Cultures (DSMZ) or the UK-based CABI. Samples from a shared sample origin from the phyllosphere remain scarce: Existing efforts have often targeted specific taxa, such as methanol-degrading methylobacteria (Knief, Frances, et al. 2010) or sphingomonads (Lundberg et al. 2022), or used an untargeted isolation approach from leaves of Northern Hemisphere plant populations (Bai et al. 2015; Reisberg et al. 2012; Qi et al. 2021; Stevens et al. 2021; Custódio et al. 2025). However, comparable resources from the Southern Hemisphere are lacking, leaving large biogeographic gaps in our understanding of plant-associated bacteria across global scales. Similarly, although bacteria are routinely isolated from leaf endophytic compartments (Hong et al. 2015), comprehensive efforts on untargeted isolation of bacterial leaf endophytes are rare (e.g. (Ramírez-Sánchez et al. 2022)) and are also lacking for *Arabidopsis* Southern Hemisphere samples.

Here, we report the isolation and full-genome sequencing of a bacterial culture collection originating from leaves of a naturalised *Arabidopsis* population in Ōtautahi (Christchurch), Aotearoa New Zealand. This region, located in the Southern Western Pacific Ocean, offers a unique opportunity to expand the global picture of *Arabidopsis*-associated microbiota. Epiphytic (leaf surface) and endophytic (apoplastic) bacteria were both isolated from individual plants, enabling direct comparison of both compartments. To assess how well the collection represents the natural community, 16S rRNA gene amplicon sequencing was performed on plant samples from the same site.

As the culture collection is, in parts, characterised and sequenced in the framework of a MSc course at Freie Universität Berlin, Berlin, Germany, the project was undertaken and completed by students in the Leaf Surface Microbiology MSc course over a four year period. Here we report on the recently achieved milestone of the full-genome sequencing of 129 isolates using Illumina sequencing and different generations of Oxford Nanopore (MinIon or Flongle) sequencing, representing 34 genera typical of the *Arabidopsis* phyllosphere, marking a key step towards the establishment of a comprehensive, genome-resolved bacterial culture collection for the *Arabidopsis* phyllosphere.

## Material and methods

### Sampling of plant material

Plant material was collected on 8th July 2020 in an urban area in Ōtautahi (Christchurch), Aotearoa New Zealand, at 43°29’57.8”S, 172°36’21.2”E, alongside the embankments of a railway line. The local *Arabidopsis* population at this location has a winter annual lifestyle, germinating in autumn (around April/May) and growing in spring. Mature, seed-bearing plants were observed at the same location in mid-October. At the time of sampling, no flowering plants were observed. Environmental conditions during collection included temperatures between 2 to 6 °C, early-morning mist followed by light rain, and a relative humidity of approximately 82%. Individual plants were sampled using sterile forceps, their roots were cut off using sterile scissors and the plants were placed in 50 mL reaction tubes. Samples were brought to the lab on ice and processed within the same days.

### Isolating bacteria from the epiphytic compartment

In the lab, five samples made up of the whole above-ground material of individual plants were carefully rinsed in sterile water to remove dust and other contaminants such as soil. Then, plant material was submerged in 10 mL phosphate buffered saline (PBS, NaCl 8 g/L, KCl 0.2 g/L, Na_2_HPO_4_ 1.44 g/L, KH_2_PO_4_ 0.24 g/L), vortexed for 30 s, sonicated for 7 min (Easy 30 H, Elmasonic), and then vortexed again for 30 s. The supernatant was then transferred into a fresh 15 mL vessel. From this supernatant, 100 µL was used to perform a serial dilution and to determine colony-forming units on Reasoner’s 2a agar (R2A, M962, HiMedia). Furthermore, aliquots were plated onto R2A and a minimal medium supplemented with methanol (1.62 g/L NH_4_Cl, 0.2 g/L MgSO_4_, 2.21 g/L K_2_HPO_4_, 1.25 g/L NaH_2_PO_4_·2H_2_O, and the following trace elements: 15 mg/L Na_2_EDTA·2H_2_O, 4.5 mg/L ZnSO_4_·7H_2_O, 0.3 mg/L CoCl_2_·6H_2_O, 1 mg/L MnCl_2_·4H_2_O, 1 mg/L H_3_BO_3_, 2.5 mg/L CaCl_2_, 0.4 mg/L of Na_2_MoO_4_·2H_2_O, 3 mg/L FeSO_4_·7H_2_O, and 0.3 mg/L CuSO_4_·5H_2_O) to isolate the bacterial colonies. The remaining supernatant was centrifuged for 10 min at 10,000 × *g* to collect the bacterial cells for DNA sequencing.

### Isolating bacteria from the endophytic compartment

After the plant material was sampled for epiphytic bacteria, its surface was sterilised by submerging the intact plant material in 70% (v/v) ethanol and vortexing for 30 s. The ethanol was discarded and the plant material was submerged in 2% (v/v) bleach and vortexed for 30 s. To remove the bleach, the samples were washed twice in PBS, then, 10 mL PBS were added and the leaves were macerated 2.5 g of 4 mm glass beads by vortexing. The resulting macerated materials were then aseptically transferred to solid media as described above. The remaining materials were used for DNA analyses.

### Long term storage of isolates

Individual colonies were restreaked at least three times to generate axenic cultures. To store the isolates, cultures were grown in liquid R2A until the culture was visually turbid. The cultures were then supplemented with 15% glycerol (v/v) and frozen at -80 °C. Isolates that did not grow in liquid culture were grown on R2A agar plates, harvested using an inoculation loop and then resuspended in PBS supplemented with glycerol (15% v/v) and frozen. All stocks were restreaked approximately one year after the initial freezing and were still viable.

### DNA extraction from environmental samples

Genomic DNA from epiphytic leaf washes was extracted with the Isolate II Genomic DNA Kit (Bioline/Meridian Bioscience, Cincinnati, USA) with slight modifications from the manufacturer’s protocol. Bacteria were ruptured in the lysis buffer GL using 0.5-mm diameter zirconium beads at 8 m/s for 30 s in a tissue lyser (Omni Bead Ruptor 24). After bead beating, 21 µL of proteinase K (22 mg/mL), 5 µL RNAse A (100 mg/mL), and 3 µL of lysozyme (300 mg/mL) were added. After the addition of lysis buffer G3, the samples were shaken at 8 m/s for 30 s in the tissue lyser. DNA of endophytic samples was extracted either with the Isolate II Genomic DNA Kit as described above or the DNeasy PowerPlant Pro Kit as recommended by the manufacturer (Qiagen, Hilden, Germany).

### Sequencing of environmental 16S rRNA gene fragments

PCR library preparation and sequencing was performed by Macrogen (Seoul, South Korea). The primers 341F (tcgtcggcagcgtcagatgtgtataagagacagCCTACGGGNGGCWGCAG) and 805R (gtctcgtgggctcggagatgtgtataagagacagGACTACHVGGGTATCTAATCC) for library preparation target the 16S rRNA gene V3–V4 region and included adapters for illumina sequencing (lower case letters) (Klindworth et al. 2013). The 16S rRNA gene amplicon sequencing was performed on the MiSeq (Illumina, San Diego, USA) platform. Approximately 100,000 paired-end reads with a length of 300 bp were generated per sample. Sequencing adapters were trimmed with trimmomatic (v0.39) (Bolger et al. 2014). Sequences were further trimmed within the DADA2 pipeline (v1.22.0) (Callahan et al. 2016), and the maximum number of ‘expected errors’ was set to 2 and 4 for forward and reverse sequences, respectively. Further, 20 bp were trimmed at the 5’ end, while 30 and 70 bp were trimmed at the 3’-end for forward and reverse sequences, respectively. Only mean Phred scores >25 per read position remained. Amplicon sequence variants (ASVs) were inferred using the DADA2 algorithm with ‘pseudo-pooling’ to increase sensitivity. Taxonomy was assigned to the sequence variants using the SILVA REF NR 99 release 138.2 dataset using DADA2. Mitochondrial and chloroplast reads were removed.

Community analyses were performed in R using the packages *phyloseq* and *vegan* (Oksanen et al. 2023; McMurdie and Holmes 2013). ASV count tables were transformed to relative abundances prior to diversity analyses. Alpha diversity was estimated using the Inverse Simpson index. Because epiphytic and endophytic communities were extracted from the sample plants, differences in alpha diversity were evaluated with a paired Student’s *t* test (α = 0.05), after confirming normality (Shapiro-Wilk test) and homogeneity of variances (Levene’s test) using the *rstatix* package (Kassambara 2023). Beta diversity was estimated based on Bray-Curtis dissimilarities on relative abundance data and visualised in ordination space by non-metric dimensional scaling (NMDS). Group centroids and dispersion were visualised with one-standard deviation ellipses at 95% confidence. Homogeneity of multivariate dispersions was assessed using betadisper (*vegan*) followed by a permutation test (n = 9,999), before differences in community composition between the epiphytic and endophytic compartments were evaluated using PERMANOVA (999 permutations).

### Genomic DNA extraction from bacterial isolates

Genomic DNA of bacterial isolates was extracted using a modified Marmur method (Salvà Serra et al. 2018). Briefly, bacteria were grown on R2A agar until colonies were well visible. Then, a loop of bacterial biomass was harvested and resuspended in 800 µL of EDTA-saline (3.723 g/L EDTA and 8.766 g/L NaCl; pH = 8.0) by vortexing in a 2 mL reaction vial. To digest the cell walls, 10 µL of lysozyme solution (300 mg/mL) was added followed by vortex mixing. Then, 7 µL of RNase A solution (10 mg/mL) was added and the sample was incubated for 37°C for 45 minutes while vortexing every 15 min. Then, 80 µL of sodium dodecyl sulfate solution (SDS, 250 g/L) was added and the sample was vortexed for a few seconds until it became viscous. The sample was then incubated at 65 °C for 10 min and vortexed once after 5 min. Then, 250 µL 5 M sodium chloride solution was added and the sample was vortexed for several seconds before 400 µL chloroform:isoamyl alcohol was added. The sample was briefly vortexed and then incubated for 15 min on an orbital shaker at 1,400 rpm. Thereafter, the sample was centrifuged for 15 min at 13,000 × *g*. The top layer of the resulting sample was transferred to a new 2 mL reaction vial while avoiding the interface between the two phases. To the new tube, 400 µL of chloroform:isoamyl alcohol was added, shaken vigorously by hand and centrifuged again for 15 min at 13,000 × *g* before the top phase was transferred to a fresh 2 mL reaction vial. This process was repeated until the interphase between aqueous and organic layer was clear. Then, for every mL of solution, 90 µL 3 M sodium acetate and 600 µL of ice cold isopropanol were added. The vial was inverted several times until DNA threads were visible. To spool the DNA, a glass rod (a glass microliter pipette that was welded shut using a bunsen burner) was used. The DNA was then dried and then resuspended in water.

### Sequencing of bacterial genomes

Due to the long term nature of this project and the integration into teaching, several sequencing technologies were used to generate bacterial full genome data. For 25 isolates, the Nextera XT DNA Library Preparation Kit was used on an Illumina NovaSeq 6000 sequencing platform, resulting in 150-bp paired-end reads. Sequencing was performed at Macrogen Oceania. A total of 101 isolates were sequenced using Oxford Nanopore Technologies (ONT) platforms with either 9.4 or 10.4.1 chemistry. Of these, 83 isolates were sequenced in-house, two isolates at Plasmidsaurus, and 16 isolates at Eurofins. Flongle sequencing was performed using the ONT Rapid Sequencing kit (SQK-RAD004) following the manufacturer’s protocol (RSE_9046_v1_revZ_14Aug2019). MinION sequencing with 9.4 chemistry was performed using the ONT Rapid Barcoding kit (SQK-RBK004) following the manufacturer’s protocol (RBK_9054_v2_revAA_14Aug2019). MinION sequencing using 10.4.1 chemistry was performed using the ONT Rapid Barcoding kit (SQK-RBK114.24) following the manufacturer’s protocol (RBK_9176_v114_revQ_27Dec2024). A complete overview of bacterial isolates identity and corresponding sequencing technologies is provided in Table S1.

### Genome assembly

Raw data received from Macrogen Oceania was processed on Curta (Bennett et al. 2020) using trimmomatic (Bolger et al. 2014) to remove adapters and low quality bases, then, genomes were assembled using SPAdes (Bankevich et al. 2012). Raw data was base-called and, where applicable, demultiplexed using the software Guppy version 6.5.7 (Oxford Nanopore Technologies). Raw data from ONT chemistry 10.4.1 was base-called and demultiplexed using the software dorado 0.9.6 (Oxford Nanopore Technologies). Data received from Plasmidsaurus and Eurofins was already base-called and hence, this step was omitted. Resulting fastq files were assembled using the software Flye using versions 2.9 up to 2.9.5 (Kolmogorov et al. 2019). Bacterial genome quality was assessed using CheckM (Parks et al. 2015) workflow “Assess Genome Quality with CheckM -v1.0.18” on the Department of Energy Systems Biology Knowledgebase (KBase) online platform (Arkin et al. 2018). Additionally, BUSCO was used using the webservice M1CR0B1AL1Z3R to assess genome completeness (Tegenfeldt et al. 2025; Shimony et al. 2025).

### Genome annotation and phylogenetic placement

Genome assemblies were annotated with Bakta v1.11.4 (Schwengers et al. 2021) using default parameters to identify coding sequences (CDS), small open reading frames (sORFs), ncRNAs, *ori* and other miscellaneous sequences. KEGG annotations were extracted to group genes by functional categories. Additional genomic features were (re-)annotated: Plasmids were detected using PLASMe v1.1 from short-read and long-read assemblies, as putative plasmid contigs, using Transformer or BLASTn (Tang et al. 2023). Only contigs were analysed that had fewer than 5% ambiguous regions, a classification score > 0.7, a length > 1 Kb, or if they were independently confirmed by BLASTn against the NCBI plasmid database. Phages were identified and annotated using PHASTEST v3.0 (Wishart et al. 2023; Phaster Arndt et al. 2016; Phast Zhou et al. 2011) using default parameters. Secondary metabolite biosynthetic gene clusters (BGCs) were detected using antiSMASH v.8.0 (Blin et al. 2025). Secretion systems were detected using TXSScan v1.1.3 (Néron et al. 2023; Abby et al. 2014; Denise et al. 2019; Abby et al. 2016). Carbohydrate-active enzymes (CAZymes) were annotated with dbCAN (Zheng et al. 2023).

Genes from the full set of strains were grouped as orthogroups using OrthoFinder v3 (Emms et al. 2025). Individual gene trees from core orthogroups present in all strains were inferred with FastTree (Price et al. 2010). A species tree was then computed from the gene trees using ASTRAL-Pro (Zhang and Mirarab 2022).

Taxonomic classification of bacterial genomes was performed with GTDB-Tk v2.5.2 with the GTDB R226 database (Parks et al. 2020; Chaumeil et al. 2022), where appropriate, taxonomy was manually curated to correct names based on ICNP. Genomes were compared by fast, alignment-free computation of whole-genome average nucleotide identity (FastANI) (Jain et al. 2018). As some genomes were nearly identical based on ANI, genomic analyses were done on 54 unique genomes (<95% ANI) and 33 representative genomes from clades with ≥95% ANI. An endophytic and epiphytic isolate from each clade were retained. Species trees and annotations were visualised using the R package *ggtree* (Yu et al. 2017).

### Genomic analysis

Phylogenetic signal and trait association with leaf compartment were assessed using phylogenetically-informed linear models and ordination. Features were grouped into genomic features (%GC, coding density, genome size, number of CDS, RNA, phage elements, and plasmid number) and functional traits (KEGG categories, BGCs, secretion systems, and CAZymes). Each feature was used as the response variable in the linear model and the leaf compartment (epiphytic vs endophytic) as explanatory variable, incorporating the species tree information inferred from orthogroups to account for phylogenetic relationships (Lajoie and Kembel 2023). For each feature, the coefficients of the regression models and the 95% CI were reported to estimate the effect size, and Pagel’s lambda (0 ≤ λ ≤ 1) to quantify phylogenetic signal, with λ ≈ 1 indicating a strong phylogenetic dependence. In parallel, a phylogenetic principal component analysis (pPCA) was performed on the root-mean-square scaled features with optimised λ to visualise multivariate trait correlations and their association with leaf environment in ordination space. Highly correlated covariates (>=90%) were filtered out. All analyses were implemented in R using *ape*, *caper*, *phytools*, and *phylolm* (Ho and Ané 2014; Paradis and Schliep 2019; Orme et al. 2023; Revell 2024).

## Results

### Endophytic and epiphytic bacterial community diversity determined by 16S rRNA gene sequencing

All communities were sequenced at a similar depth, with read numbers of 64,620 ± 20,019 and 35,222 ± 9,530 (mean ± SD) for endophytic and epiphytic samples, respectively (Figure S1). However, the endophytic communities were heavily enriched in ASVs classified as mitochondrial and chloroplast rRNA sequences which were filtered out prior to further analyses. After filtering, three endophytic samples had a total number of reads <65, while the remaining had 289 and 933 reads (Figure S1).

Taxonomy was assigned using the SILVA database (REF NR99 release 138.2). The taxonomic distribution (Class level) showed that Alphaproteobacteria were dominant taxa in both endophytic (39.1%) and epiphytic compartments (77%), followed by Actinomycetes (25% and 10.5%, respectively) (Figure 1A). In some cases, Gammaproteobacteria were dominant over Alphaproteobacteria in the endophytic compartment. Certain groups were only found in one compartment: Bacilli (27.4%), Nitrospiria (3.1%), and Planctomycetia (1.0%) were exclusively detected in the endophytic compartment, and Terriglobia (3.8%), Cyanobacteriia (6.6%), and Deinococci (1.4%) occurred only in the epiphytic fraction. Of the 43 ASVs that were detected endophytically, 13 overlapped with the epiphytic compartment (Figure S2). Alpha diversity was significantly higher in epiphytic than in endophytic communities (Figure 1B, t-test on Inverse Simpson indexes, *t*(4) = 4.9, *p* = 0.008), and based on Bray-Curtis dissimilarities, community composition also differed between compartments (Figure 1C, PERMANOVA, *R*^2^ = 0.22, *p* = 0.028).

**Figure 1.**
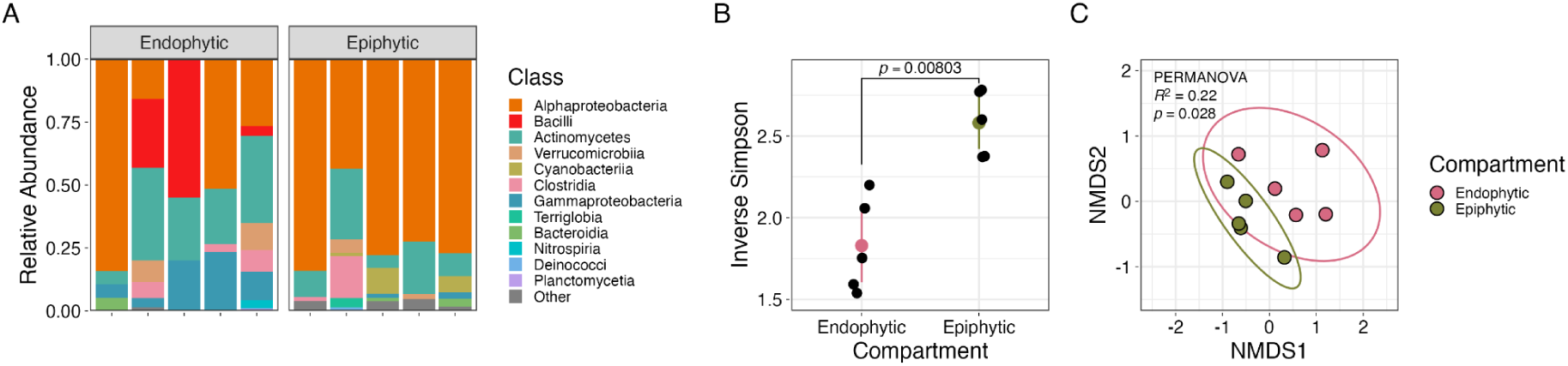
Bacterial community structures on natural *Arabidopsis thaliana* from Aotearoa New Zealand. Environmental DNA was extracted from the above ground epiphytic and endophytic compartments of five plants, and the 16S rRNA gene was targeted for amplicon sequencing. (A) Relative abundances of abundant bacterial classes across leaf compartments. “Other” includes ASVs representing <1% of the total community. (B) Alpha diversity estimated by the Inverse Simpson index. Mean and 95% CI of each group are indicated in colour. (C) Community composition based on Bray-Curtis dissimilarities. *R*^2^ and *p*-value based on PERMANOVA on communities grouped by leaf compartment (endophytic and epiphytic).

### Collection of isolates

A total of 799 bacterial strains were isolated, including 499 from epiphytic samples, of which 100 were picked from minimal media supplemented with methanol, and 300 from endophytic samples (Table 1). All isolates were photographed after they formed visible colonies on R2A agar plates (Schlechter et al. 2025). Additional isolates, indicated in brackets, originated from samples that contained more than one bacterial species.

**Table 1.** Bacterial strains isolated, stocked, and sequenced from *Arabidopsis thaliana* leaf material from Ōtautahi Christchurch (Aotearoa New Zealand)

| Plant sample | Epiphytic isolates R2A selection |  |  | Epiphytic isolates methanol selection |  |  | Endophytic isolates R2A selection |  |  |
| --- | --- | --- | --- | --- | --- | --- | --- | --- | --- |
|  | Picked | Stocked | Sequenced | Picked | Stocked | Sequenced | Picked | Stocked | Sequenced |
| #1 | 100 | 54 | 18 | 20 | 15 | 0 | 60 | 30 | 4 |
| #2 | 100 | 75 | 22 | 20 | 18 | 1 | 60 | 27 | 13 |
| #3 | 100 | 59 | 16(+1) | 20 | 20 | 1 | 60 | 39 | 11(+1) |
| #4 | 51 | 46 | 10 | 20 | 20 | 2 | 60 | 38 | 13 |
| #5 | 51 | 47 | 5(+1) | 20 | 20 | 0 | 60 | 39 | 10 |
| **Total** | 402 | 281 | 73 | 100 | 93 | 4 | 300 | 173 | 52 |

Numbers in parenthesis represent strains that were recovered from mixed genome samples.

### Genome completeness and resolution of mixed assemblies

A total number 50 endophytic and 74 epiphytic isolates were successfully sequenced. Their genomes were annotated and checked for completeness and contamination using CheckM (Table S2). While most assemblies were predicted to have low contaminations (0–3.44%), three assemblies were outliers: AC348, AC405, and AC640 showed predicted contaminations of 98.28–100%. Upon closer inspection, it was apparent that these cultures were not axenic. These genomes making up these non-axenic assemblies were assessed for quality and separated into individual genomes; the assembly of AC348 was obtained using ONT sequencing and contains two circular contigs: contig 1 (5,206,036 bp, 49× coverage) was most similar to *Xanthomonas* spp., while contig 2 (5,181,883 bp, 91× coverage) was closest to *Pseudomonas* spp. The resulting assemblies were separated and renamed AC348a and AC348b, respectively. The assembly of AC405 was obtained using ONT sequencing and consisted of 3 contigs. Contigs 1 and 2 were assigned to *Pseudomonas* spp., with 16× coverage, while contig 3 matched *Chryseobacterium* (26× coverage). These assemblies were renamed AC405a and AC405b, respectively. The assembly of AC640 was obtained using Illumina sequencing and consisted of 173 contigs. To separate mixed genomes, MaxBin2 (v2.2.4) was used (Wu et al. 2016), resulting in two bins: “bin 1” (22 contigs, >300× coverage) assigned to *Luteimonas*, and and “bin 2” (44 contigs, >15× coverage) assigned to *Agrococcus*. The resulting assemblies were renamed AC640a and AC640b, respectively.

After separating the non-axenic genomes, a total number of 129 genomes were analysed, CheckM predicted completeness values for all genomes ranging from 18.75% to 100%, with 121 genomes above 90% completeness (Table S2). Genome completeness according to BUSCO, ranged from 85.5% to 100%, with four sequences showing a completeness below 95% (Table S2).

### Phylogenetic placement of the sequenced isolates

We sequenced and assembled 129 bacterial genomes from the *Arabidopsis* phyllosphere from a natural population in Aotearoa New Zealand and placed them into a phylogenetic tree (Figure 2). Most isolates belonged to the phylum Pseudomonadota (68.2%), followed by Actinomycetota (18.4%), Bacteroidota (10.9%), and Bacillota (2.3%) (Figure 3). We have yet to cultivate representatives from the phyla Acidobacteriota, Cyanobacteriota, Desulfobacterota, Nitrospirota, or Verrucomicrobiota detected via 16S rRNA gene amplicon sequencing (Figures 1-3) in these *Arabidopsis* samples. The relative distribution of taxa can be found

**Figure 2.**
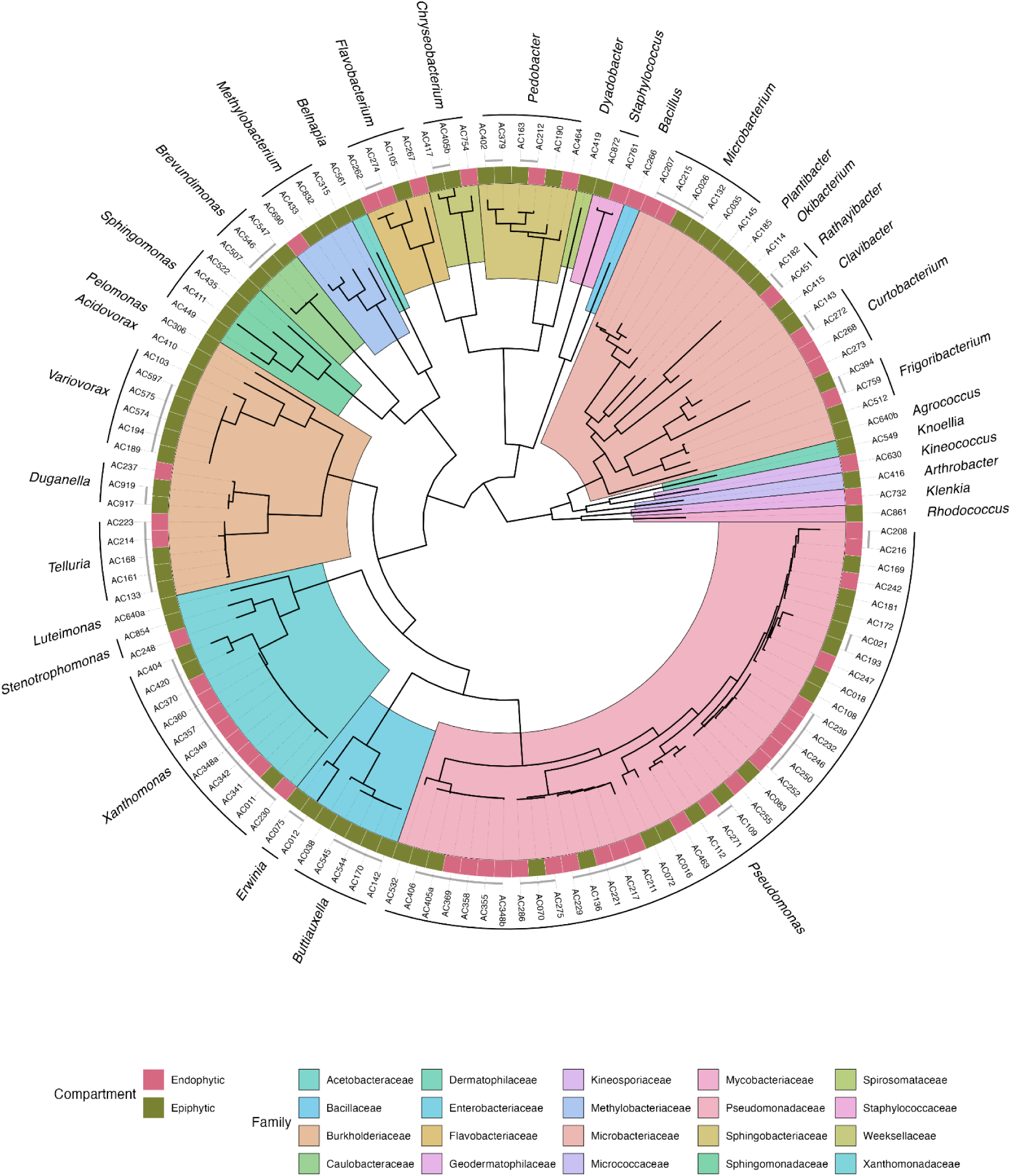
Phylogenetic tree of the AC collection. Maximum-likelihood species tree reconstructed by OrthoFinder from single-copy orthogroups. Taxon names correspond to the strains included in this study. Outer ring colors denote the compartment of isolation (endophytic vs. epiphytic). Gray lines indicate groups of strains sharing ≥95% average nucleotide identity (ANI). Classification was undertaken using GTDB-Tk. Clades indicate genera.

**Figure 3.**
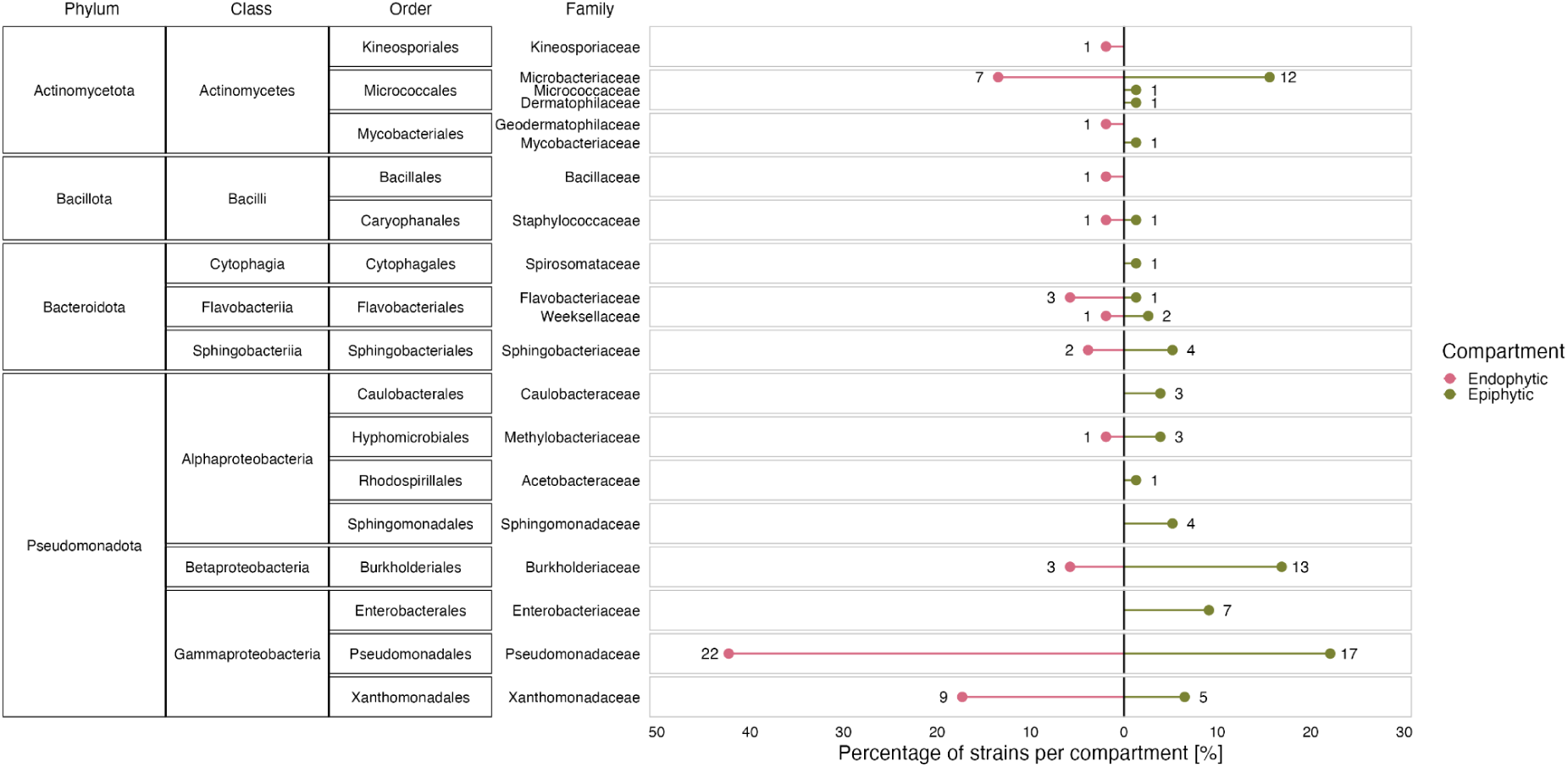
Taxon representation within the full genome sequenced isolates. Percentage of taxon represented within each leaf compartment. Number indicates the number of isolates.

Within the detected phyla, the collection comprises the following eight classes: Gammaproteobacteria (46.5%), Actinomycetes (18.6%), Betaproteobacteria (12.4%), Alphaproteobacteria (9.3%), Flavobacteriia (5.4%), Sphingobacteriia (4.7%), Bacilli (2.3%), and Cytophagia (0.8%). These classes represent 16 orders: Pseudomonadales (30.2%), Micrococcales (16.3%), Burkholderiales (12.4%), Xanthomonadales (syn. Lysobacterales, 10.9%), Enterobacterales (5.4%), Flavobacteriales (5.4%), Sphingobacteriales (4.7%), Hyphomicrobiales (syn. Rhizobiales, 3.1%), Sphingomonadales (3.1%), Caulobacterales (2.3%), Caryophanales (syn. Staphylococcales, 1.6%), Mycobacteriales (1.6%), Kineosporiales (0.8%), Cytophagales (0.8%), Rhodospirillales (syn. Acetobacterales, 0.8%), and Bacillales (2.3%).

In total, the dataset contains 20 families: Pseudomonadaceae (30.2%), Microbacteriaceae (14.7%), Burkholderiaceae (12.4%), Xanthomonadaceae (syn. Lysobacteraceae, 10.9%), Enterobacteriaceae (5.43%), Sphingobacteriaceae (4.7%), Beijerinckiaceae (3.1%), Flavobacteriaceae (3.1%), Sphingomonadaceae (3.1%), Caulobacteraceae (2.3%), Weeksellaceae (2.3%), Staphylococcaceae (1.6%), Acetobacteraceae (0.8%), Bacillaceae (0.8%), Dermatophilaceae (0.8%), Geodermatophilaceae (0.8%), Kineosporiaceae (0.8%), Micrococcaceae (0.8%), Mycobacteriaceae (0.8%), Spirosomataceae (0.8%).

Several isolates clustered with known leaf-associated bacteria: 17 with *Pseudomonas koreensis*, four with *Pseudomonas fluorescens*, and two with *Pseudomonas syringae* (AC016 and AC072), both of which caused disease symptoms in *Arabidopsis* in infection assays undertaken as part of the *Leaf Surface Microbiology* graduate class studies course (data not shown). Similarly, the isolate AC341, which is closely related to the phytopathogen *Xanthomonas dyei*, causes lesions on *Arabidopsis* leaves *in vitro* (data not shown).

Across the endophytic and epiphytic isolates, most taxa overlapped between leaf environments, particularly members of Pseudomonadales, Microbacteriaceae, Xanthomonadales, Burkholderiales, and Sphingobacteriaceae. However, certain groups were unique to each environment (Figure 3). Isolates of Geodermatophilaceae, Kineosporiaceae, Bacillaceae were only detected in the endophytic environment so far, while in the epiphytic environment Enterobacteriaceae, Sphingomonadaceae, Caulobacteraceae isolates were detected.

### Genomic features of the AC collection

Of the 129 isolates, 54 strains were unique (ANI <95%), while the remaining strains were grouped into ANI clusters (Figure S3A, ≥95% ANI). Representative members were then selected based on genome completeness and contamination, and, when applicable, at least one isolate from each compartment was included (Figure S3B). In total, 87 genomes were retained for further genome comparison of two sub-collections: 30 from the endophytic compartment and 57 from the epiphytic compartment.

We compared structural genomic features, such as genome size, GC content, number of CDS and different types of RNAs, among others (Table S3); and mobile elements, such as phages (Table S4) and plasmids (Table S5), between leaf compartments while accounting for phylogeny. A strong phylogenetic signal was observed in genome structure (median Pagel’s λ = 1.0, Figure 4A), which indicated that closely related strains present similar genomic traits. The presence and structure of phages had a much weaker signal (median Pagel’s λ = 0.53), while the number of plasmids had a Pagel’s λ of 0.94.

**Figure 4.**
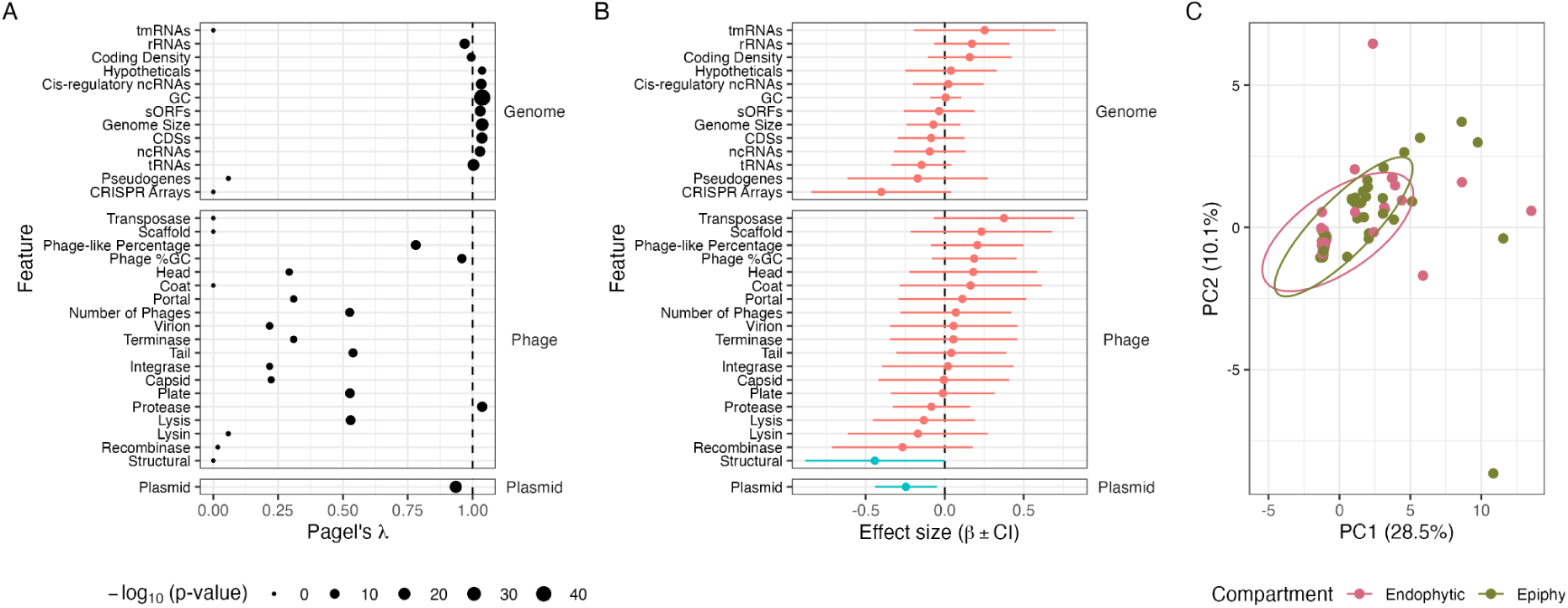
Phylogenetic structure and effect of leaf compartment on genomic structures of the AC collection. (A) Phylogenetic signal (Pagel’s λ) for structural and mobile element features. Values close to 1 indicate strong conservation along the tree. (B) Estimated effects (β) and 95 % confidence intervals for structural and mobile element features obtained from phylogenetic generalised linear models (phylolm). Positive values indicate higher trait values in epiphytic strains relative to endophytic strains; red: *p*-value > 0.05; blue: *p*-value < 0.05. (C) Phylogenetic principal component analysis (pPCA) of scaled genomic structural features. Points are coloured by compartment (epiphytic vs. endophytic); ellipses show 95% confidence intervals.

Phylogenetic regression showed that the endophytic vs epiphytic compartment showed little to no difference in explaining the variation of genome structure among the isolates (Figure 4B). In most cases, the effect sizes for nearly all genomic features were close to zero between endophytic and epiphytic strains (*p* > 0.05). The exceptions were the presence of structural phage protein-coding genes (β = -0.44, *p* = 0.049) and number of plasmids (β = -0.25, *p* = 0.014), which in both cases were higher among the endophytic strains. However, these traits explained little of the variation (*R*^2^ = 3.3–5.7%). Consistent with the regression analysis, a phylogenetic principal component analysis (pPCA) explained the 38.6% of the variation of the genomic features in the first two principal components, however, it showed no clear separation of endophytic and epiphytic strains (Figure 4C). Together, these results indicate that the genomic structure of our collection is shaped primarily by phylogeny, with little evidence of convergence associated with the leaf compartment.

### Functional features of the AC collection genomes

We functionally annotated and compared the genomes of each representative isolate (*N* = 87) using KEGG categories for general metabolic pathways (Table S6), secretion systems (Table S7), secondary metabolism (Table S8, biosynthetic gene clusters, BGCs), and carbohydrate-active enzyme-coding gene families (Table S9, CAZymes). Overall, most of the broad functional groups analysed showed a high phylogenetic signal (Figure 5A, Pagel’s λ > 0.82, *p* < 0.05, FDR correction). However, individual functional categories differed depending on their type. Most variation was found in secretion systems, with T3SS showing the lowest phylogenetic signal (median Pagel’s λ = 0.171), followed by T4SS and T6SS. Next, BGCs associated with non-ribosomal peptide synthetases (NRPS, λ = 0.53) and terpenes (λ = 0.66) were less determined by phylogeny than others. Within KEGG groups, glycan biosynthesis (λ = 0.68) and metabolism and cellular community (biofilm, λ = 0.82) had the lowest phylogenetic signal.

**Figure 5.**
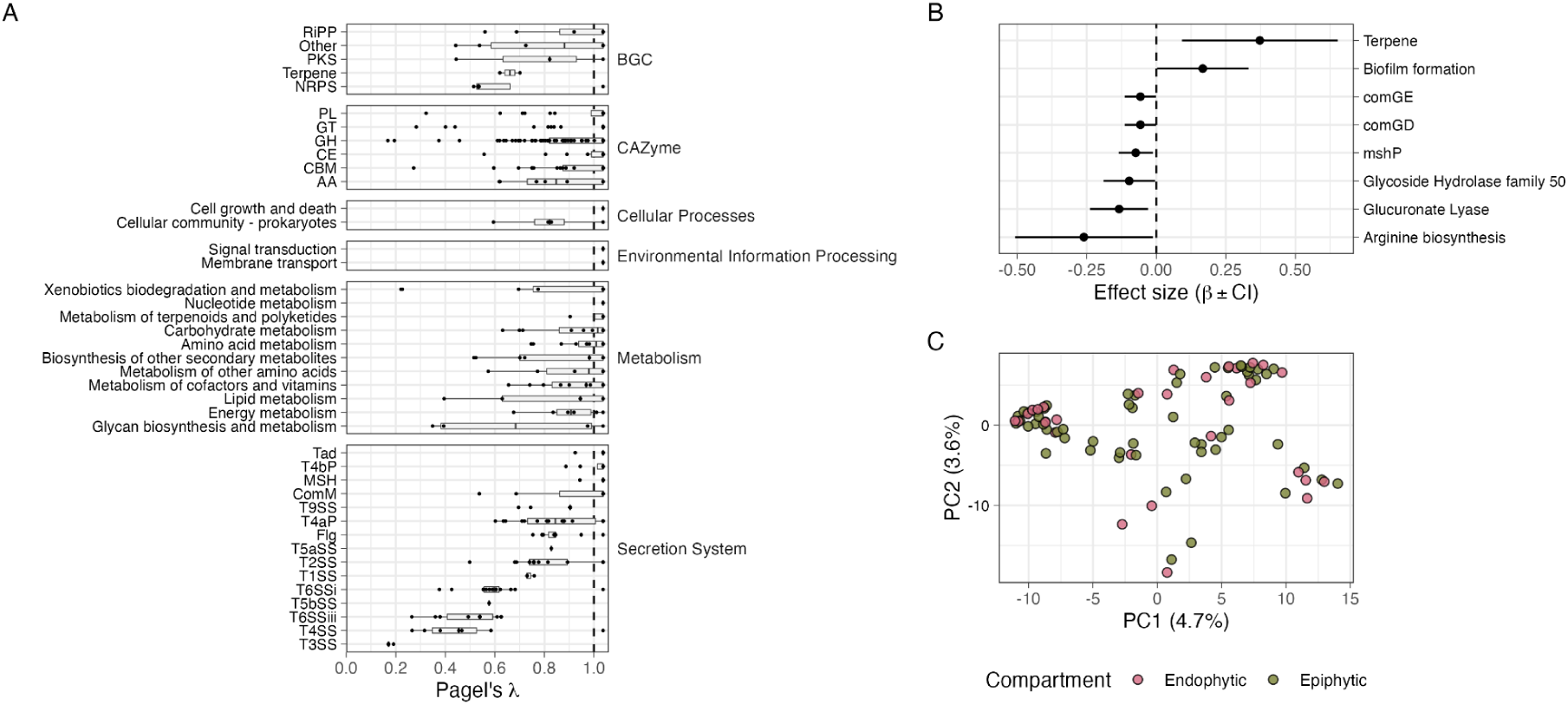
Phylogenetic structure and effect of leaf compartment on functional features within the AC collection. (A) Phylogenetic signal (Pagel’s λ) for functional groups associated with biosynthetic gene clusters (BGC), carbohydrate-associated enzymes (CAZymes), KEGG categories (Metabolism, Environmental Information Processing, and Cellular Processes), and secretion systems. Values of λ close to 1 indicate strong phylogenetic conservation along the tree as opposed to the isolation source. (B) Estimated effects (β) between isolation source (endophytic vs epiphytic) and 95 % confidence intervals for functional features obtained from phylogenetic generalised linear models (phylolm), with *p*-value < 0.05. Positive values indicate higher trait values in epiphytic strains relative to endophytic strains. (C) Principal component analysis (PCA) of scaled functional features. Points are coloured by compartment (epiphytic vs. endophytic).

Phylogenetic regression showed only a few functions with significant differences between endophytic and epiphytic isolates (Figure 5B). Among the traits tested, terpene biosynthetic clusters (β = 0.37, *p* = 0.0099) and biofilm formation (β = 0.167, *p* = 0.046) were enriched in epiphytic isolates. Terpene biosynthetic clusters were present in 24 epiphytic strains, mostly from Microbacteriaceae, Sphingobacteriaceae, Pseudomonadaceae, and Burkholderiaceae, in contrast to only six endophytic strains. Biofilm formation genes were represented in epiphytic strains belonging to Sphingomonadaceae, Pseudomonadaceae, Enterobacteriaceae, Burkholderiaceae, and Staphylococcaceae.

By contrast, endophytic genomes were enriched in genes associated with arginine biosynthesis (β = -0.26, *p* = 0.040), mostly from Pseudomonadota and Actinomycetota genomes. A much smaller effect was observed for the CAZymes glucuronate lyase (PL27, β = -0.13, *p* = 0.012), enriched in endophytic *Pseudomonas* and *Pedobacter*, and glycoside hydrolase (GH50, β = -0.097, *p* = 0.040), present in endophytic Actinomycetota and Bacteroidota. Additionally, competence-related genes (*comGE*, β = -0.074, *p* = 0.019; and *comGD*, β = -0.057, *p* = 0.046) and a mannose-sensitive hemagglutinin pilus gene (*mshP*, β = -0.074, *p* = 0.019) were enriched endophytically, mainly driven by Bacillota and Pseudomonadota. An ordination of all functional categories by principal component analysis showed no clear separation between endophytic and epiphytic genomes, and only 8.3% of the variance explained by the first two components (Figure 5C). Genomes clustered primarily by phylogeny rather than by leaf compartment. Overall, these results are consistent with the structural genome analyses: functional gene content is strongly determined by phylogeny, with only limited and scattered signatures of convergence associated with leaf compartment.

## Discussion

### Bacterial endo- and epiphytic diversity

Leaves are colonised by recurring bacterial communities, with few alternative dominant genera (Lundberg et al. 2022), similar to human intestinal enterotypes (Arumugam et al. 2011). Reflecting this observation, the bacterial populations from *Arabidopsis* specimens sampled in Ōtautahi (Christchurch), were dominated by sphingomonads and pseudomonads and were consistent with previous observations that these two groups frequently dominate *A. thaliana* phyllosphere communities (Lundberg et al. 2022). While the culture collection was dominated by pseudomonads, potentially due to their higher culturability, the overall diversity patterns of the collection reflected those observed from the environmental samples, that is, the average Inverse Simpson indices at the genus level were comparable for isolates versus environmental samples (epiphytic: 2.28 vs 2.06; endophytic: 1.98 vs 1.70, respectively).

Our 16S rRNA amplicon sequencing and isolate-based approaches show that epiphytic communities are richer than endophytic communities, which is consistent with a majority of similar studies (Mahmoudi et al. 2024; Agler et al. 2016; Sanjenbam and Agashe 2024). This outcome is expected, given that the endophytic compartment is more immigration-limited compared to leaf surfaces, as entry into the apoplast requires passage through barriers such as the cuticle and stomata, and successful evasion of host immune responses, thereby restricting colonisation opportunities (Smets et al. 2023).

Overall, both the 16S rRNA amplicon survey and the bacterial isolate collection represented bacterial taxa commonly detected and isolated from *Arabidopsis* and or other plant leaves, including *Sphingomonas*, *Methylobacterium*, *Pseudomonas*, *Flavobacteria*, *Pedobacter*, *Rhodococcus* (Bai et al. 2015; Aydogan et al. 2018; Agler et al. 2016; Almario et al. 2022). Additionally, the taxonomy of many isolates resemble taxa from a previous *Arabidopsis* isolation survey, the “LSPHERE” collection (Bai et al. 2015)(Genomes in Figure S4 whose epithet starts with ”Leaf”). This demonstrates that bacterial communities and represented taxa on *Arabidopsis* are similar across broad geographic distances, suggesting that the phyllosphere environment and the host plant species acts as an important selection filter to shape community composition (Schlechter et al. 2019; Redford et al. 2010; Karasov et al. 2024).

Unexpectedly, several species that were found in the strain collection were not detected during NGS. This can, in part, be explained by the relatively low sequencing depth of the endophytic samples, and differences in lysis efficiency between taxa during metagenomic DNA isolation compared to the ease of culturability (Willner et al. 2012). In particular, the paucity of betaproteobacterial taxa detected in the 6S rRNA gene amplicon datasets stood out in our study, and is consistent with previous reports of inefficient lysis of this group (Willner et al. 2012). Similar discrepancies between culture-dependent and culture-independent datasets have been highlighted in recent synthetic community (SynCom) research, reflecting persistent taxonomic blind spots across both approaches (Northen et al. 2024). These mismatches can result from uneven amplification efficiency, incomplete reference databases, and the underrepresentation of certain lineages in molecular surveys. Establishing culture collections in parallel with community profiling has therefore been emphasised as essential to capture the full ecological and functional diversity of plant-associated microbiota and to design SynComs that are taxonomically representative (Northen et al. 2024). Integrating genomic, phenotypic, and ecological data from isolates will help determine how well the cultured strains reflect the natural bacterial community associated with plants and reduce biases introduced by primer selection and extraction efficiency.

A clear pattern in leaf compartment-specific representation was observed and, while this might be a result of undersampling of the different collections, it does evidence fundamental differences in the communities and selection of particular members. Especially Comamonadaceae, Sphingomonaceae and Enterobacteriales were represented only among epiphytic isolates, while most other taxa were represented in both collections, including some nearly identical genotypes. The latter indicates a regular exchange between the epiphytic and endophytic environment. Although it is unclear which compartment acts as the source and which one as sink, the epiphytic compartment is more likely to act as a primary source, as it harbors larger population densities, is generally richer, and has fewer migration barriers than the endophytic compartment.

### Genomic characterisation of endophytic and epiphytic culture collections

Overall, the obtained genomes were generally of good quality with little contaminations (Table S2), which serves as a good foundation to utilise the genomes for future genomic-driven studies (Northen et al. 2024; Jing et al. 2024).

Our comparative analysis highlights that genomes of leaf-associated bacteria are shaped mostly by phylogeny rather than adaptations to the leaf environments. Across our dataset, most structural genomic traits, including genome size, GC content, coding density, as well as functional categories showed substantial phylogenetic signals, consistent with previous work showing that bacterial traits tend to be conserved (Martiny et al. 2013; Goberna and Verdú 2016; Tamames et al. 2016). This suggests that the ability of bacteria to colonise either the leaf surface or the apoplast largely reflects inherited traits rather than extensive genomic specialisation to each leaf compartment.

Differences in genomic structures between epiphytic and endophytic strains are less likely to be due to specialisation of mutualistic symbionts in *Arabidopsis*, but more likely indicates activity of transposable elements and accumulation of other repetitive DNA in the endophytic compartment (Manzano-Marín and Latorre 2016). As expected, mobile genetic elements and horizontally transmitted elements showed weaker phylogenetic signals, reflecting how horizontal gene transfer decouples bacterial traits from vertical inheritance. Secretion systems such as T3SS, T4SS, and T6SS showed low phylogenetic signal, consistent with these elements being mobile across taxa (Frank et al. 2005) and highly divergent (Wallner et al. 2021; Gillespie et al. 2015). Similarly, BGCs displayed low phylogenetic structure and are often plasmid-borne, particularly in host-associated bacteria (Saati-Santamaría 2023), and they also rapidly evolve through frequent rearrangements and recombination events (Medema et al. 2014; Chase et al. 2021). Glycan biosynthesis and biofilm formation similarly stood out as two metabolic pathways with low phylogenetic signal. Glycan modification and biofilm formation could be involved in mediating interactions with other microbes and being recognised by the plant host (Amador et al. 2025; Wanke et al. 2021), suggesting that traits may also be influenced by ecological pressures in the leaf environment.

Biofilm formation-related and terpene biosynthesis genes were enriched in epiphytic isolates, indicating that the role of those genes are more important during epiphytic colonisation. Biofilm formation on leaves is an important trait of microbial colonisers of leaves (Morris and Monier 2003) and likely allows bacterial adhesion to leaves to avoid removal by rain events (Schlechter et al. 2019; Barthlott and Neinhuis 1997). Bacterial and plant terpene metabolism has been investigated in the context of the chemical crosstalk between plants and the microbial colonisers (Junker and Tholl 2013). Plant-produced terpenes have been proposed to act as bactericidal compounds that reduce the number of bacteria that successfully colonise leaf surfaces (Gao et al. 2005).

Conversely in the endophytic compartment, genes relating to the utilisation of arginine and CAZymes glucuronate lyase were enriched with a noteworthy effect size. The utilisation of arginine by epiphytic has been detected on leaves before, but has so far not been linked to the endophytic compartment (Ryffel et al. 2016). Our finding suggests that arginine utilisation is possibly more important in the endophytic compared to the epiphytic compartment. The overrepresentation of glucuronate lyase is likely linked to the degradation of glucuronic acid components of plant cell wall pectin polymers (Hugouvieux-Cotte-Pattat et al. 2014).

### Future use of the strain collection

The collection of full genome sequenced bacteria will facilitate the construction of representative microbial communities that are purpose-tailored to study the impact of microbial communities on plant functioning and microbial ecology of the phyllosphere and multitrophic interactions of plant-microbe-herbivore interactions (Müller et al. in prep). Beyond serving as a taxonomically defined resource, culture collections provide a foundation for assembling experimentally tractable SynComs that can be used to investigate colonization dynamics, priority effects, and microbial interactions under controlled gnotobiotic conditions (Vorholt et al. 2017). Comparable approaches have been successfully applied in *Arabidopsis* and other plant models to dissect the mechanisms underlying host specificity, community assembly, and microbe–microbe interactions in root-associated microbiota (e.g., (Wippel et al. 2021)). However, such defined-community experiments remain rare for phyllosphere systems, where the environmental constraints and interaction networks differ from those in soil and the rhizosphere (e.g. (Carlström et al. 2019; Bai et al. 2015)). Establishing and extending existing resources for leaf epi and endophytic-associated bacteria therefore fills an important gap and enables the experimental dissection of microbial interactions in an aboveground habitat.

The availability of fully sequenced isolates with defined ecological origins further enhances the reproducibility and comparability of plant–microbe interaction studies. By enabling researchers to reassemble natural-like bacterial consortia *in vitro* or *in planta*, this resource bridges the gap between descriptive microbiome studies and mechanistic experiments. Such standardized strain collections are therefore essential for moving toward reproducible model systems that allow the direct testing of hypotheses about microbial function, host adaptation, and community resilience.

Our culture collection and genomic information support ongoing efforts towards standardisation in plant–bacteria interaction research. Our dataset aligns with initiatives to develop standardised microbial resources that serve as reference communities and ensure the reproducibility of gnotobiotic experiments (Northen et al. 2024). Establishing and characterising such collections contributes to broader goals of harmonising experimental protocols, generating benchmark data, and promoting the open sharing of strains and sequence information. Ultimately, these resources will not only support fundamental research but also provide a foundation for applied studies aiming to manipulate leaf-associated microbiota for enhanced plant health and sustainability.

## Data availability

Genome sequencing results can be found in the EMBL-ENI ENA BioProject accession PRJEB98369. Additional sequences will be made available in the BioProject as the collection expands. Images of each strain and data sets associated with this study are available in Zenodo (https://doi.org/10.5281/zenodo.17812155) (Schlechter et al. 2025). The R scripts used for data analysis are available in the GitHub repository https://github.com/relab-fuberlin/AC_collection.

## Acknowledgements

We acknowledge Ngāi Tūāhuriri as mana whenua of the rohe in which these microorganisms were collected. We further acknowledge that we used the high performance computer cluster CURTA at Freie Universität Berlin. This work was supported by a Marsden Grant (UOC1704) to MRE.

## Author contributions

MMi sampling, culture maintenance, DNA extraction, Illumina sequencing, data analysis; RS conception, sampling, MinION sequencing, culture maintenance, data analysis, data organisation, writing-original draft; RJ sampling, culture maintenance; CW culture maintenance, DNA extraction; SO culture maintenance; CS sampling, culture maintenance; MMa culture maintenance, DNA isolation; EK culture maintenance, data organisation; SH culture maintenance, DNA isolation; MO culture maintenance, DNA isolation; LPV writing-original draft; MBS conception, writing-editing; MRE funding acquisition, conception, sampling, culture maintenance, Illumina sequencing, MinION sequencing, data analysis, writing-original draft. Students of the course Leaf surface microbiology were involved in initial laboratory characterisation and sequencing/ sequence assembly verification, due to their alumni status, it was not possible to communicate with them. All other authors have read and approved the submitted manuscript.

