## Supplemental figures for "Comparative genomics of epiphytic and endophytic bacterial culture collections from *Arabidopsis thaliana* in Ōtautahi (Christchurch), Aotearoa New Zealand"

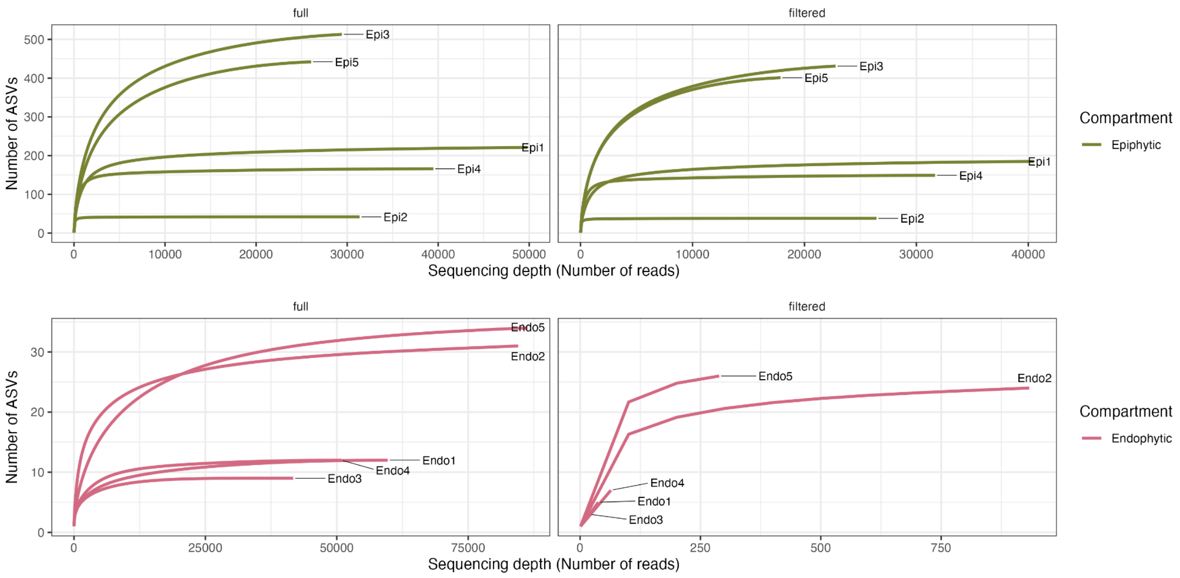


**Figure S1.** Rarefaction curves before (left) and after (right) chloroplast and mitochondrial ASVs were filtered out.

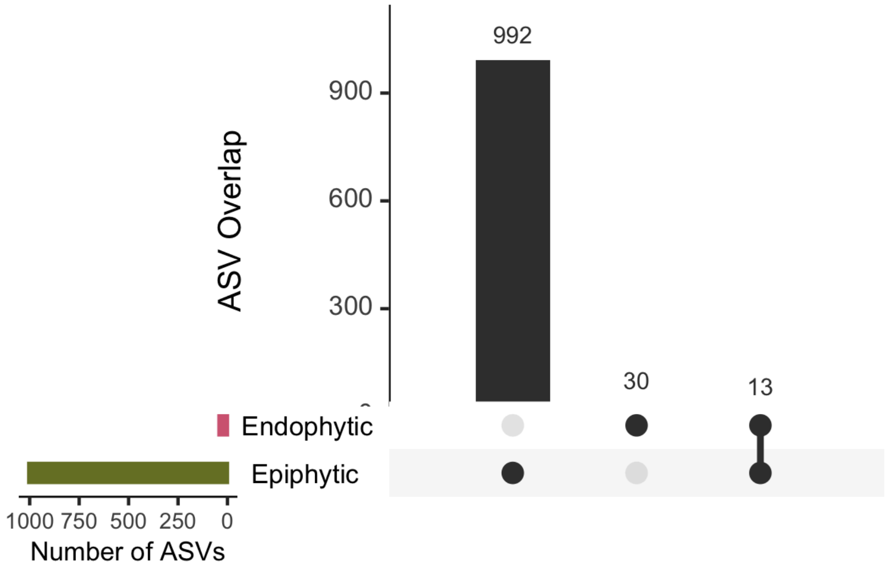


**Figure S2.** Upset plot of overlapping ASV in epiphytic and endophytic samples.

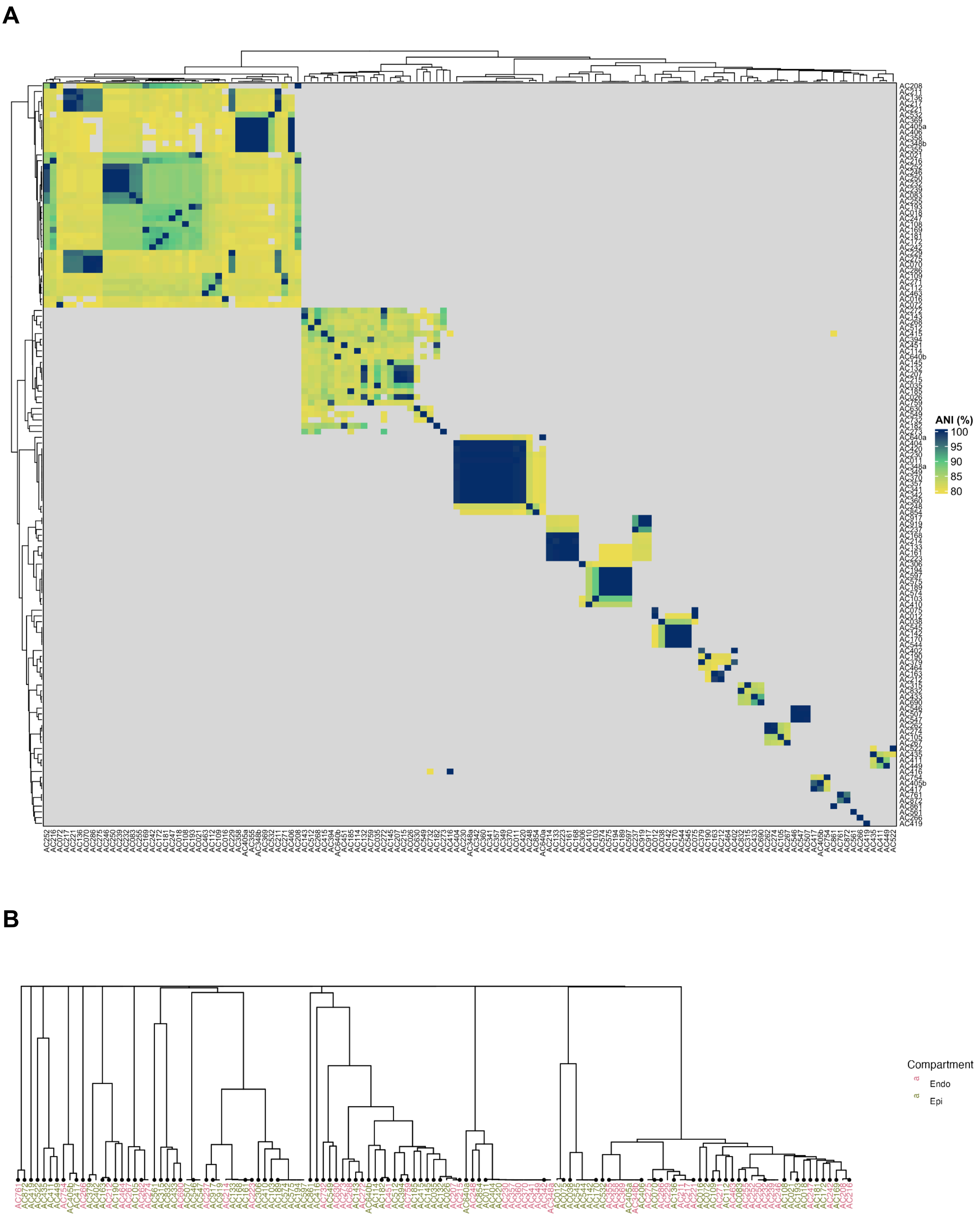


**Figure S3. FastANI. (A)** Heatmap showing the pairwise ANI similarity (%) between isolates. **(B)** Dendrogram based on ANI similarities. Dots indicate strains that have been selected for analysis, with one representative (based on CheckM contamination and completeness) for clades with ANI > 95% and compartment (endophytic or epiphytic).

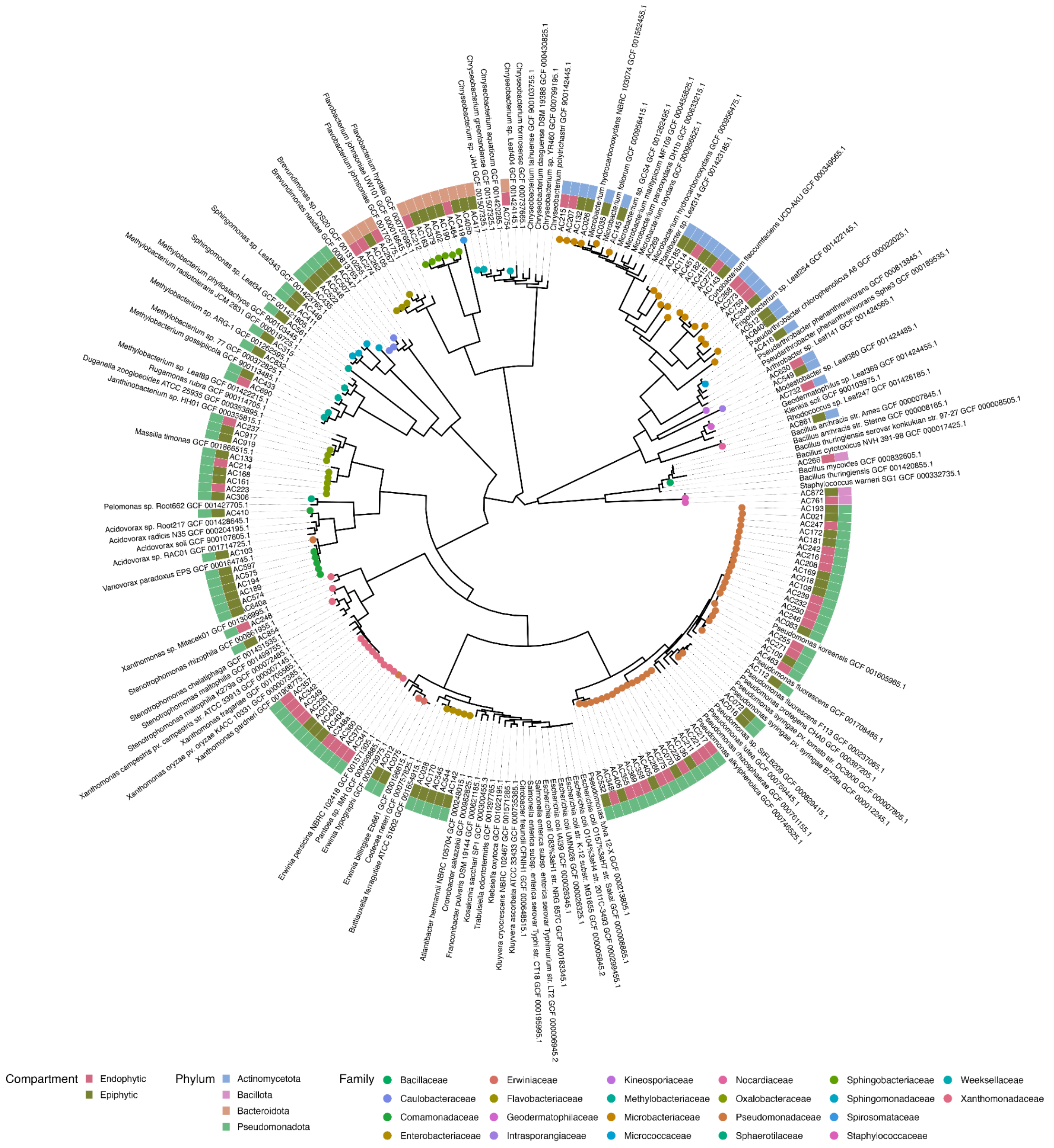


**Figure S4.** Phylogenetic tree of the isolates generated by multilocus sequence analysis of 49 concatenated core genes. The tree was produced using the Build Microbial SpeciesTree (v1.6.0) workflow on Kbase.org, additionally to the isolates sequenced in this work, closely reference sequences were automatically added to provide phylogenetic and taxonomic context to the isolates. Isolate origins are highlighted in the outermost circle, phylum in the second circle and families of the isolates are highlighted as colored dots at the end of the branches.
